# Ocular speech tracking persists in blindness, but its dynamics and oculo-cerebral connectivity depend on visual status

**DOI:** 10.1101/2025.10.21.682842

**Authors:** Kaja Rosa Benz, Larissa Reitinger, Fabian Schmidt, Davide Bottari, Anne Hauswald, Olivier Collignon, Nathan Weisz

## Abstract

While eye movements have been shown to track the speech envelope, it is unknown whether this reflects a hard-wired mechanism or one shaped by (lifetime) audiovisual experience. Further, questions remain about whether ocular tracking is modulated by speech intelligibility and which brain regions drive these synchronized eye movements. Here, we investigate ocular speech tracking in blindfolded early blind, late blind, and sighted individuals using magnetoencephalography (MEG) and source-reconstructed oculomotor signals while participants listened to narrative speech of varying intelligibility. We found that oculomotor activity tracks acoustic speech features and, unlike neural speech tracking, is not modulated by intelligibility. Interestingly, we found effects reflected in two frequency-specific components: a low-frequency (∼1 Hz) effect present across all groups, indicating that visual experience is not required, and a high-frequency (∼6 Hz) effect reduced in early- and late-blind individuals. Moreover, this finding is not driven by cerebro-ocular connectivity, as late-blind individuals exhibit stronger connectivity between the eyes and the left temporal cortices without a corresponding increase in ocular tracking. In conclusion, ocular speech tracking seems to respond selectively to acoustic but not to intelligibility features of speech, and it does not require visual experience to develop. It may thus represent a hard-wired oculomotor mechanism within the oculo-cerebral network involved in speech processing.

**Significance Statement:** Eye movements provide a unique window into the interaction between auditory and visual systems. By studying early blind, late blind, and sighted individuals, we demonstrate that speech-related eye movements arise from at least two distinct mechanisms: a low-frequency component that occurs independently of (lifetime) visual experience and is linked to processing of acoustic speech features, and a high-frequency component shaped by prior visual exposure. Importantly, speech intelligibility - unlike its impact on neural measures - does not modulate these ocular responses. This dissociation suggests that eye movements reflect mechanisms of spoken language processing that are independent of intelligibility, thereby revealing novel pathways of auditory-motor coupling and broadening our understanding of sensory integration in the absence of vision.

## Introduction

In most settings, speech perception is an audiovisual process, drawing on a hierarchy of processing stages to support comprehension [33, 3, 24, 47]. These processes range from sensory encoding of acoustic speech features to higher-level integration of linguistic features [27, 28]. Efficient processing relies on neural systems dynamically tracking these features [17, 44], known as neural speech tracking [43]. Recent findings demonstrate that speech is also tracked by eye movements. Informed by a substantial body of research on audiovisual integration highlighting the involvement of eye movements in speech processing [11, 32, 37, 38, 46], Gehmacher et al. [20] established that eye movements track the speech envelope of continuous, narrative, attended speech – a phenomenon termed ocular speech tracking. Ocular tracking was strongest for the attended speaker in multi-speaker settings, and higher tracking was observed in participants reporting lower comprehension [20, 51], suggesting a compensatory attentional role under demanding listening conditions [51]. Listening effort is often probed by degrading intelligibility using vocoded speech, which reduces spectral detail while preserving the temporal envelope [53]. Our previous work shows that neural speech tracking modulates as a function of speech intelligibility [15, 26, 50], but whether ocular tracking shows a similar modulation or reflects sensitivity to acoustic dynamics remains unknown. A central open question is whether ocular speech tracking requires concurrent retinal input. Previous work tested only sighted participants with open eyes but no task-relevant visual input, leaving unresolved whether ocular tracking persists without visual input. Another question concerns whether ocular tracking develops in the absence of (lifetime) visual experience. If the development of ocular tracking requires visual experience, early-blind individuals should lack it; if it reflects a hard-wired mechanism, it should occur independently of visual status. We previously showed that during silent lip-reading, auditory deprivation is influencing ocular tracking of the (unheard) speech envelope, as late deafs exhibit increased ocular tracking while early deafs did not show envelope tracking [5]. Although controlling for eye movements did not explain neural effects in that work, in auditory speech bidirectional interactions between oculomotor and neural systems [20, 40] appear to play an important role in ocular speech tracking. Establishing how ocular activity couples with the brain during listening to speech could clarify whether eye movements are driven by top-down cortical control or contribute bottom-up signals to auditory processing in the absence of visual input. Early-blind individuals retain functional circuitry between motor cortex and oculomotor activity [35]. Generally, early-blind individuals are known to have superior hearing abilities compared to sighted individuals, which is associated with activation in the visual cortex [49], which is also involved in the processing of spoken words [41]. This leads to the hypothesis that, in the absence of vision, eye movements may still support auditory processing. However, findings of reduced eye-visual cortex connectivity in early blindness [35] challenge this idea. Alternatively, it is possible that the eyes of blind individuals are still involved in speech perception but relay information directly to the ears [39, 34], without involving the visual cortex. Since these eye-to-ear connections do not rely on visual input [1], and blind participants exhibit eye-movement patterns comparable to those of sighted individuals [25], ocular speech tracking might occur independently of being blind or not. Here, we directly address these open questions by reanalyzing magnetoencephalography (MEG) data from a previously published dataset (early blind, n= 17; sighted, n= 14) [55] extended by a late blind group (n= 16). Participants were blindfolded and listened to continuous, narrative speech presented at three intelligibility levels (natural, 8-channel vocoded, 1-channel vocoded). We extracted eye-movement time-series data using source reconstruction techniques and computed coherence to investigate ocular and neural speech tracking and functional oculo-cerebral connectivity. Specifically, we asked: (1) Does ocular speech tracking occur while participants listen to speech in the absence of concurrent visual input, and is it modulated by intelligibility? (2) Does its presence and strength depend on lifetime visual experience? (3) Which neural systems drive ocular tracking, and how does oculocerebral connectivity vary with visual status? By addressing these questions, we test whether ocular speech tracking reflects a hard-wired, experience-independent mechanism or one shaped by visual experience, and investigate how it relates to established neural speech tracking processes.

## Materials and Methods

### Participants

Seventeen early blind (7 male; 33.6 ± 10.55 years; 20-67 years), 16 late blind (9 male; 45.5 ± 16.62 years; 26-71 years), and 14 sighted participants (4 male; 33.2 ± 13.06 years; 20-63 years) participated in the present study. Groups were matched in terms of age and biological sex [55]. All participants were proficient braille readers and native Italian speakers. None of them suffered from a known neurological or peripheral auditory disease. All early blind participants were either totally blind or severely visually impaired from birth; however, two reported residual visual perception before the age of 3, one before the age of 4, and one participant lost their sight completely at age 10. Residual visual perception of diffuse light was present in 8 early blind participants. Causes of vision loss included retinal damage or detachment (10), optic nerve damage (3), infection of the eyes (1), microphthalmia (2), and hypoxia (1). In the late blind group, the age of blindness onset varied from childhood to adulthood. Residual visual perception of diffuse light was present in 10 late blind participants, and 5 had no residual vision. Causes of late blindness included retinopathy (7), congenital glaucoma (3), damaged optic nerve (4), eye injury (1), and congenital cataracts (1). Participants signed informed consent forms before participation. The project was approved by the Ethics Committee of the University of Trento and was in accordance with the Declaration of Helsinki.

### Experimental design

The present study builds on data from Van Ackeren et al. (2018) [55]. Auditory stimuli were presented within the magnetically shielded MEG room via stereo loudspeakers employing a Panphonics Sound Shower Two amplifier, calibrated to a consistent and comfortable volume level across all participants. The presentation of the stimulus was controlled using Psychophysics Toolbox 3 (http://psychtoolbox.org; RRID:SCR002881) running in MATLAB (RRID:SCR001622), executed on a Dell Alienware Aurora workstation operating on 64-bit Windows 7. To eliminate visual input, both sighted and blind individuals were blindfolded throughout the experimental session. The room was dimly illuminated to permit video-based observation of participants during MEG acquisition. Experimental instructions were provided via pre-recorded audio messages spoken by one of the experimenters. The speakers mean syllable rate was 5.99 Hz (std: 3.01 Hz) [calculated with syllable nuclei [16] computed using the Python library parselmouth [31]]. The stimulus set consisted of 14 one-minute excerpts derived from well-known Italian audiobooks (i.e., Pippi Longstocking and Candide). Two additional conditions were generated through channel vocoding using Praat software [8]. Specifically, the original audio signal was decomposed into either one (1-channel) or eight (8-channel) logarithmically spaced frequency bands. For each band, the amplitude envelope was extracted and used to amplitude-modulate band-limited Gaussian noise (noise carrier). The modulated noise bands were then recombined into a single audio signal, preserving the overall temporal envelope while progressively degrading spectral detail. This manipulation preserved the overall amplitude envelope while gradually degrading spectral detail. In perceptual terms, the 8-channel version produced substantial vocal distortion without impairing intelligibility, whereas the 1-channel version rendered speech entirely unintelligible. In total, 42 stimuli were presented across seven pseudo-randomized blocks. Each block included two exemplars from each condition, with no repetition of narrative content within a block. Representative audio samples are provided in the supplementary materials. To assess comprehension, each stimulus segment was followed by a single declarative sentence summarizing the story content. Participants were instructed to attend closely to each passage and to indicate whether the statement was true or false using a non-magnetic response box, operated with the index and middle fingers of the right hand.

### MEG Data Recording

The experiment was conducted at the MEG Lab at the Center for Mind/Brain Sciences (Laboratorio di magnetoencefalografia at CiMeC) in Rovereto in Italy. Whole-head MEG was acquired using a triple-sensor (204 planar gradiometers; 102 magnetometers) 306-channel system (Elekta Neuromag, Helsinki, Finland). Continuous recordings had a sampling rate of 1000 Hz and were band-pass filtered online between 0.1 and 300 Hz. Individual head shapes were digitized with a Polhemus FASTRAK 3D tracking system (Polhemus, Colchester, Vermont, United States). Five localization coils were used to continuously record the head position of the subject.

### MEG Data Preprocessing

Data preprocessing was performed in MNE-Python [22]. Initially, all recordings were processed with a signal space separation algorithm (Maxwell filter; [54]) that removes external magnetic interference and corrects for within-session head movement. For this, the data from each individual participant were realigned to their mean head position across blocks. For independent component analysis (fast ICA; [29]), a 1 Hz high-pass filter was applied to the raw data and it was downsampled to 150 Hz, yielding 50 components. Cardiac artifacts were identified automatically with the default function in MNE Python. The components were applied to the original raw data, and the cardiac artifact removed. This ICA-corrected data was filtered between 0.1 and 16 Hz using a Blackman filter and the default MNE raw filter settings and also downsampled to 150 Hz. We subsequently attempted to identify consistent ocular-related ICA components using semiautomated methods, including correlation with frontal channel activity, template matching [12], and evaluation of component topography symmetry. However, as participants were blindfolded with a heavy bandage to avoid voluntary eye movements and blinks, we were not successful in consistently identifying them across groups (no convincing component for 17% of the participants). Given the inadequacy of ICA to model ocular activity in this particular study, we implemented a source reconstruction approach for this purpose. Note that for the sensor plots and statistics we used the magnetometers only, but for the source reconstruction all channels were included.

#### Source projection of MEG data

We employed a spatial filtering approach (LCMV beamformer) to reconstruct both ocular and neural sources. Source projection of the data was performed with MNE-Python [22]. An automatic coregistration pipeline was used to align the FreeSurfer “fsaverage” template brain [18] to each participant’s head shape. After an initial fit using the three fiducial landmarks, the coregistration was refined with the iterative closest point algorithm [6]. Head-shape points located more than 5 mm away from the scalp were automatically omitted. Since a single-layer boundary element model (BEM; [2]) does not include the eyes, a sphere model was used as the BEM parameter in the present study. Next, a surface source space was defined using the ico-4 subdivision (5124 sources in total, 2562 per hemisphere), and the eyeballs were added as a volume source space with 514 sources. Based on the fsaverage template, the eyes were modeled as two spheres positioned at the locations of the eyes in the template MRI. The forward operator (i.e., lead field matrix) was then computed using the individual coregistration, the sphere model, and the mixed source space (surface for the brain and volume for the eyes). For further analysis, the principal component of each region from a multi-modal parcellation of the human cerebral cortex (HCPMMP1; [21]) was used for visualization and statistical analyses. Consistent with the approach for brain regions, the first principal component from the PCA of the eye-source reconstructions was used for further analysis.

#### Speech envelope extraction

Amplitude envelopes of the auditory stimuli were derived using the Chimera toolbox [14] in combination with custom scripts, following the methodology outlined by Gross et al. (2013) [23]. First, the audio files were band-pass filtered into nine frequency bands from 100 to 1000 Hz using a fourth-order Butterworth filter. To avoid phase shifts relative to the original signal, filtering was applied in both forward and backward directions. Frequency bands were evenly spaced to correspond with the human basilar membrane’s tonotopic organization. The analytic amplitude for each filtered segment was computed as the absolute of the Hilbert transform. Subsequently, the amplitude envelopes across all bands were summed and normalized to a maximum value of one. This composite envelope was then also downsampled to 150 Hz and temporally aligned with the MEG recordings and processed using identical analysis pipelines. For the control envelope, the envelope of the respective audio file was cut into two pieces and the two parts were exchanged.

### Speech tracking

To investigate neural speech tracking, two complementary analytical approaches are commonly used: frequency-domain measures, such as coherence, and regression-based methods, such as the multivariate temporal response function (mTRF) [36]. Both approaches have also previously been applied to investigate ocular speech tracking [20, 51, 5]. To obtain an integrated picture with both methods, we decided to utilize and present both approaches with coherence and mTRFs. As demonstrated in previous studies, the results have yielded either consistent or complimentary results across varying levels of intelligibility [15] and diverse linguistic backgrounds and exposure [45]. The coherence results are presented in the main body of the manuscript, and the (mainly consistent) mTRF results are illustrated in the supplemental material.

#### Coherence

For the coherence analysis, the data were segmented into epochs of 6 seconds to ensure sufficient resolution of the low-frequency oscillations. A constant detrending procedure was applied to the epochs, thereby removing any offset without altering the linear trend. In addition, baseline correction was applied using the interval from 0 to 0.5 seconds, ensuring that subsequent analyses were referenced to this period. Coherence was then calculated using the spectral-connectivity-epochs function from MNE-Python, applied to the epoched data. We used multitaper spectral estimation, without adaptive weighting, and without frequency averaging, to preserve spectral resolution. Coherence was computed across the frequency range between 0.1 Hz to 8 Hz to include delta (*<*4 Hz) and theta (4-8 Hz) bands. The resulting measures are cerebro-acoustic coherence as a measure for neural tracking, and ocular-acoustic coherence as a measure for ocular tracking. To test the connectivity between the eyes and the brain, we used the imaginary part of coherence, in order to avoid volume conduction effects [42]. All other settings were identical to the coherence calculation. Imaginary coherence yields positive and negative values, with the sign indicative of the directionality of connectivity (here: positive indicates oculo-cerebral (i.e., eye-to-brain) connectivity, and negative cerebro-ocular (i.e., brain-to-eye) connectivity). These were analyzed in a descriptive manner. For statistics, we used the absolute value of the imaginary coherence.

#### Multivariate Temporal Response Function

For the estimation of mTRFs, we used Eelbrain’s boosting implementation [9] of the full timeseries data. For each of the three conditions, we concatenated all blocks, resulting in 8 minutes of data per participant, on which the boosting analysis was ran. As [7] suggest 5-15 seconds length of the partition, the data was partitioned into 50 folds (leading to around 10 s per fold), with one fold used as the test set in each iteration. The model was fit for a temporal window from -0.4 to 0.8 s relative to stimulus time series (see [51]). We used an l1 error metric to promote sparsity in the model. The mTRFs were estimated using a basis function width of 50 ms, allowing temporal smoothing of the response functions. To avoid overfitting, selective stopping was enabled, so that boosting was halted independently for each predictor when it no longer improved model performance on the validation data.

#### Statistics

To test for differences in frequency bands, time series, and sensors, we ran a cluster-based permutation test to correct for multiple comparisons in eelbrain [10]. For this, the whole range was used: all magnetometers, full frequency range for the coherence (0.25 to 8 Hz), and a time range of interest for the mTRF(−100 to 800 ms). For the sensor plots in 1b) we pre-selected the sensors with the highest 10 percent of envelope coherence at 1Hz, and called them auditory sensors in order to make sure that we are not confusing auditory activity with the ocular activity data. When comparing for the control vs. the actual envelope, a cluster-based permutation t-test was conducted, while for comparing the groups and conditions a cluster-based permutation ANOVA (groups*conditions) was run. For the source reconstructed data, we conducted a cluster-based permutation test with controlling for neighboring regions. We plot and plot the uncorrected F-values in Fig. 2b), thereby enabling interpretation of the spatial patterns at 1 and 6 Hz, and show the resulting clusters of the cluster based permutation test in Fig. 2c).

**Figure 1:**
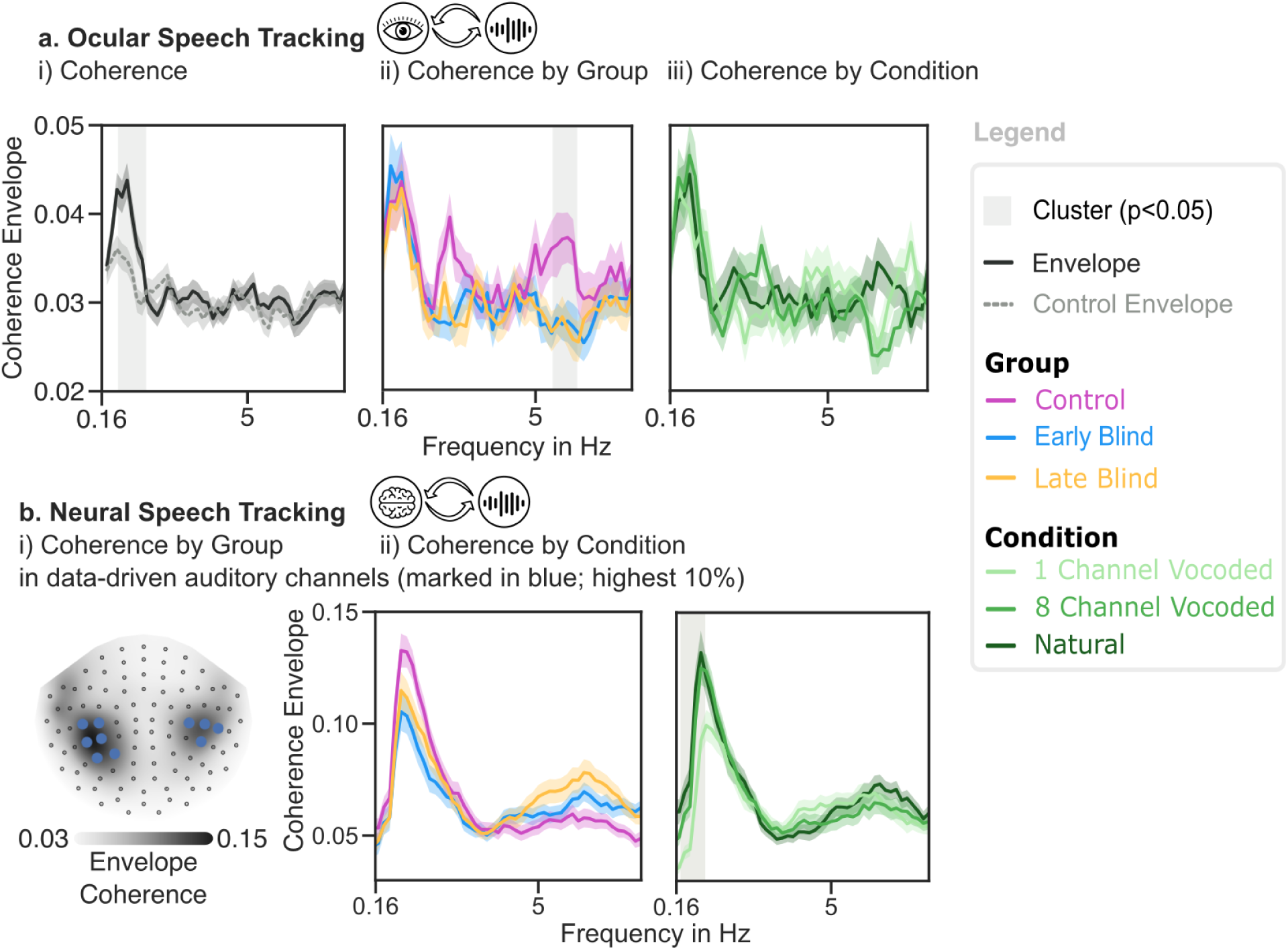
a) Ocular response to speech is i) present with eyes closed ii) increased in sighted individuals from 5.5-6.33 Hz compared to the early and late-blind individuals iii) not modulated by intelligibility. b) Topography of the envelope coherence at 1 Hz was used to select auditory channels, to show if the ocular tracking is influenced by the envelope tracking in auditory channels.

**Figure 2:**
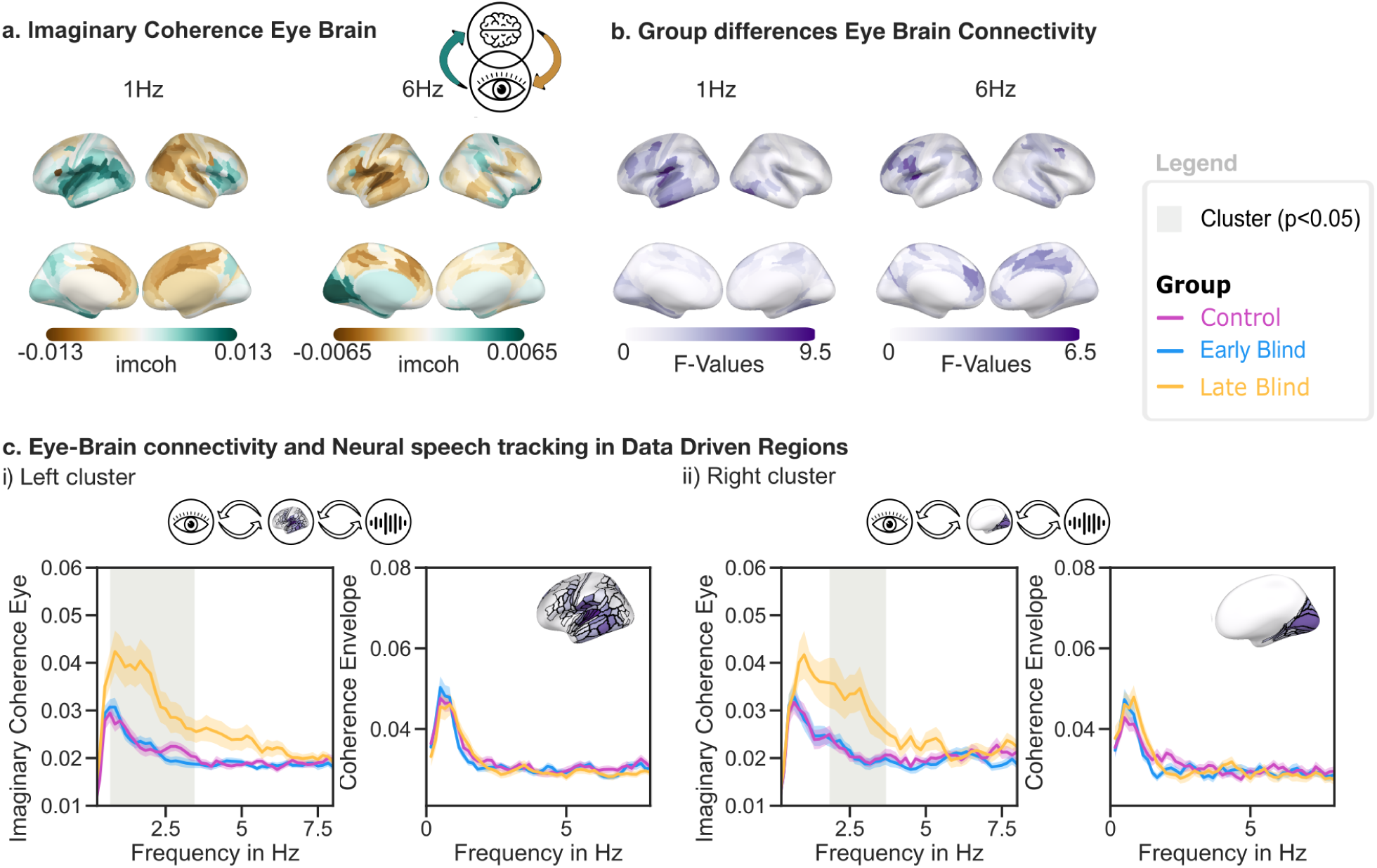
a) Imaginary coherence without any masking. The imaginary part of coherence indicates oculo-cerebral connectivity when positive (green) and cerebro-ocular connectivity when negative (brown). b) Group comparison: uncorrected F-values of the absolute value of the imaginary coherence. c) Brain areas extracted from the cluster based permutation test of the absolute value of the imaginary coherence. These plots reveal significantly increased oculo-cerebral imaginary coherence in the late-blind group. Follow-up analyses of cerebro-acoustic coherence in these regions showed no group differences.

## Results

In the present study, we investigated ocular and neural speech tracking in early blind, late blind, and sighted individuals while they blindfoldedly listened to continuous, narrative speech presented at three different intelligibility levels (natural, 1-channel vocoded, 8-channel vocoded) without relying on direct eye-tracking methods. Instead, we used source reconstruction techniques to estimate ocular activity. Additionally, we investigated the connectivity between ocular and neural sources depending on visual status. Our primary aim was to assess whether (lifetime) visual input is needed for ocular speech tracking to occur, to investigate whether ocular tracking modulates with speech intelligibility, and to depict which neural systems drive ocular tracking.

### Ocular activity during eyes-closed and blindfolded listening tracks speech envelope

In a first step, we show across all groups that the reconstructed eye movements track speech acoustics, even when listening with closed and blindfolded eyes. This means that oculomotor activity tracks speech without depending on changing retinal input elicited by eye movements.

We compared tracking of the acoustic envelope of the presented audios to a control envelope and found higher tracking of the actually presented acoustic envelope. This was the case for coherence in low frequencies from 0.5 to 1.5 Hz (*t*(47) = 50.34, *p < .*005, see Fig. 1.a.i) and for the reconstruction accuracy (*t*(47) = 4.27, *p < .*001, 1 − *β* = .99, see Supp. ia)). For the mTRF weights, three significant time windows were identified: (1) 220–273 ms (*t*(47) = 85.87, *p < .*001), (2) 280–487 ms (*t*(47) = 280.92, *p < .*001) and (3) 500–800 ms (*t*(47) = 341.15, *p < .*001, see Supp. i.b).

### Ocular speech tracking is not related to intelligibility

To further investigate the functional properties of ocular speech tracking, we tested whether oculomotor activity is modulated as a function of speech intelligibility. We propose that if there is a condition effect such that the acoustics of intelligible speech are tracked more strongly than those of unintelligible speech (i.e., 1-channel vocoded), then closed-eye ocular activity would be responsive to the intelligibility of speech. Conversely, if not, we suggest that closed-eye ocular activity tracks speech envelope modulations irrespective of intelligibility. In line with the latter, no significant clusters were found in the coherence effects for oculo-acoustic coherence (see Fig. 1.a.iii) or in mTRF reconstruction accuracies. Moreover, there was no significant main effect of condition (*F* (44, 2) = 0.73, *p* = .485, *η*_*g*_^2^ = 0.009).

To rule out the possibility that ocular effects reflect volume conduction from auditory sources, and to validate our approach against prior evidence showing that neural speech tracking is modulated by intelligibility [55, 26, 15, 50], we compared the neural data across the three intelligibility conditions. Both coherence and mTRF analyses confirmed the directionality of the previously reported results. Selecting the sensors with the strongest response, in order to avoid measuring this auditory effects also in the eyes, we find a neural condition effect from 0.33 to 1.17 Hz compared to the (*F* (44, 2) = 46.99, *p <* 0.05). A similar pattern was also observed for the mTRF approach (See Supplements). Descriptively, neural tracking was strongest in the natural condition and weakest in the 1-channel vocoded condition, with the 8-channel vocoded condition falling in between. This graded pattern replicates previous evidence [15, 50] showing that cortical entrainment to speech acoustics is enhanced by intelligibility. This pattern is not present in the ocular data, suggesting that ocular tracking is reflective of a different mechanism of the perceptual system than neural tracking.

### Ocular tracking of acoustics changes as a function of visual experience

As the mechanisms underlying oculomotor tracking of speech acoustics remain unclear, we compared eye movements in early blind, late blind, and sighted participants to assess whether this phenomenon depends on visual experience or instead reflects hard-wired, experience-independent mechanisms. Coherence analysis revealed a significant main effect of group in the 5.5–6.33 Hz frequency range (*F* (44, 2) = 23.95, *p* = .014; see Fig. 1.a.ii), with sighted participants showing stronger ocular speech tracking than both blind groups. Importantly, at ∼1 Hz - the frequency at which we observed a tracking effect in the previous step (see Fig. 1.a.i) - no group differences were found. This pattern suggests the involvement of two distinct underlying mechanisms: one at ∼1 Hz, present regardless of visual experience or sightedness, and another one at ∼6 Hz, observed only in sighted individuals. For the mTRF reconstruction accuracies, no significant main effect of group (*F* (44, 2) = 0.27, *p* = .765, *η*_*g*_^2^ = 0.006) was observed (see Supp. i.c). To ensure that the neural and ocular effects are distinct, in line with the condition comparison, we plotted the F-value topography of the neural speech tracking group comparison at 1 Hz and 6 Hz and selected the sensors with p-values *<* 0.01, to plot the full coherence spectrum. Strikingly, these patterns differ from those observed for ocular tracking: around 1 Hz, the post-hoc tests show increased tracking in the control group between 0.50 and 1.83 Hz (*F* (44, 2) = 41.90, *p* = .038), whereas around 6 Hz, the late blind group shows enhanced tracking between 5.33 and 7.16 Hz (*F* (44, 2) = 66.58, *p* = .006) (see Fig. 1.b.i). Note that these effects are expected to occur, as we pre-selected the significant channels. We present them here only to compare the patterns to the ocular effects, and they should not be interpreted otherwise.

### Strongest oculo-cerebral connectivity patterns in late-blind individuals

At ∼1 Hz, ocular tracking was strongest compared to the control envelope, and at ∼6 Hz, the group difference was strongest. Therefore, we investigated the connectivity between ocular and neural activity at both frequencies of interest (1 Hz and 6 Hz). Using imaginary coherence, across the two frequencies of interest, we observed distinct patterns of oculo-cerebral connectivity: at 1 Hz, left temporal and inferior frontal regions appeared to follow the eyes, whereas at 6 Hz, the left temporal and inferior frontal regions preceded the eyes. To compare the three groups in this measure, we used the absolute value of the imaginary coherence, focusing on the strength rather than the directionality of the connectivity, and ran an ANOVA. F-values across all parcellated areas for the frequencies where we found the ocular effects are illustrated in Fig. 2.b. The cluster-based permutation tests comparing the oculo-cerebral connectivity reveals 2 widespread clusters: on the left side, it starts at 0.167 Hz and ends at 3.33 Hz (*F* (44, 2) = 3187.8, *p <* 0.001) including many regions (See Fig.2.c.i and for a list Supp.2), peaking at the left secondary auditory cortex (L-PBelt-ROI-lh). On the right hemisphere the cluster peaks occipitaly at the V7 (R-V7-ROI-rh) the significant difference is a bit slower: from 2 until 3.6 Hz (*F* (44, 2) = 608.07, *p <* 0.01). See all regions in (See Fig.2.c.ii and for a list look at Supp.2)

For these regions, we plotted the the full frequency range (0.16–8 Hz) in Fig.2.c.i and Fig.2.c.ii (left panel) and marked the previously on whole-brain level encountered cluster in grey. Finally, to test whether the enhanced oculo-cerebral connectivity or the neural tracking activity is driving the ocular tracking of the envelope, we compared neural envelope coherence across groups within the data-driven regions. No significant group differences were observed (Fig. 1.v.a,b, right panel). This shows that the stronger oculo-cerebral connectivity observed in the late blind group was not leading to stronger neural speech tracking.

## Discussion

Here, we present the first study on whether visual input is needed for the oculomotor system to track incoming auditory information, to characterize how oculomotor activity changes as a function of speech intelligibility, and to investigate oculo-cerebral connectivity to explore which neural systems drive ocular tracking. We compared ocular and neural tracking in early blind, late blind, and sighted individuals while they listened to continuous, narrative speech. Our results show that closed eyes track the speech envelope, reflected in a low-frequency (∼1 Hz) effect across all groups and a high-frequency (∼6 Hz) effect only in sighted individuals. We suggest this pattern to reflect two different processes of ocular speech tracking - a hard-wired low-frequency process that is independent of visual experience and a high-frequency one only employed in a functionally intact visual system. In contrast to neural tracking, which has previously been shown to modulate with speech intelligibility [26, 50, 15], ocular tracking does not. Thus, we propose ocular speech tracking to reflect a functionally different mechanism of speech processing than neural speech tracking.

### Hard-wired low-frequency ocular tracking

We show that low-frequency (∼1 Hz) oculomotor activity in early blind, late blind, and sighted individuals tracks the speech envelope, even in the absence of visual input, with closed eyes (see Fig. 1.a.i; 1 Hz peak). This shows that ocular speech tracking is not dependent on changing retinal input which would be the result of moving the eyes when they are open. The finding that ocular speech tracking is present across all groups additionally suggests that it does not occur as a consequence of previous exposure to audiovisual speech, for example, by interacting with a speaker. Whether ocular speech tracking is hard-wired or develops in response to audiovisual speech exposure was answered in part by one of our previous studies [5]. There, we demonstrated enhanced ocular tracking in late-deaf compared to hearing individuals so stronger visual involvement in late deafs (as the group is relying more on lip movements) might be increasing ocular tracking. Its absence in early-deaf individuals facing visual speech (lip movements), suggests that it needs auditory exposure to speech to develop. Complementing this, the present results suggest that while auditory input is necessary for the emergence of ocular tracking, visual input is not – challenging the idea that ocular speech tracking depends on audiovisual learning and instead supporting the notion of a hard-wired mechanism involved in auditory processing. This slow tracking of speech peaking at 1 Hz is primarily involved in segmenting speech without periodic activity based on speech onsets, such as the beginning of a sentence [13]. It could be also reflecting intonation units, that are reflected in the envelope and generate a stable low frequency rhythm at 1 Hz [30]. Investigating oculo-cerebro activity, instead of a strong occipital involvement, we show left-temporal oculo-cerebral connectivity at ∼1 Hz (see Fig. 2.a). This suggests that slow stimulus-locked eye movements support speech processing in auditory regions. We find that activity in the medial frontal and parietal regions, which are suggested to drive attentional processes, precedes ocular activity. Interestingly, this falls in line with previous research proposing ocular speech tracking to be attention-driven [20, 51]. Further, ocular activity precedes activity in left-temporal regions, suggesting an eye-ear connection that enables direct auditory information transmission [39]. Following this line of reasoning, the low-frequency effect seems to be driven by attentional processes and possibly an internal model of speech, which would explain why auditory [5] but not visual experience is required. Further research is needed to gain a more comprehensive understanding of the (functional) role of this low-frequency effect.

### High-frequency ocular tracking develops in response to previous exposure to audiovisual speech

As discussed above, a central question of the present study was whether ocular speech tracking is a hard-wired phenomenon or shaped by visual experience. Our results suggest that both factors play a role. Although the three groups do not differ in the low-frequency (∼1 Hz) effect, sighted controls show increased tracking in the high-frequency (∼6 Hz) effect compared to the blind groups. The speech feature supported at ∼6 Hz is the syllable rate, which might be reflected in this effect. The enhanced tracking in the sighted group suggests that a general engagement of the visual system influences ocular speech tracking, even when blindfolded. Syllabic transition rates expressed in lip movements and speech might be learned by audiovisual experience. Since sighted individuals integrate audiovisual information when listening to a speaker in everyday life, it could make sense that auditory input is sent to the eyes. Intriguingly, in this data the mean syllable rate is at 6 Hz so low-level auditory processing would happen in temporal regions for local computations and would drive the ocular tracking of speech. Consequently, greater involvement of the visual system predicts stronger phase locking of eye movements to the speech envelope. Coherent with previous research suggesting bidirectional oculo-cerebral connectivity [40], we find that a left-temporal network precedes ocular activity at ∼6 Hz, indicating that these processes may enhance ocular tracking at this frequency in sighted individuals. In summary, at a fast pace (∼6 Hz), the ocular system is employed to process auditory information, peaking exactly at the syllable rate of the heard speech, which is only the case for sighted individuals.

### Oculomotor activity of blindfolded individuals tracks speech acoustics independently of intelligibility

In line with previous research suggesting that ocular speech tracking reflects an attentional mechanism supporting challenging listening situations, posing the idea that listening to vocoded speech might enhance ocular speech tracking [5, 20, 51], we expected to find ocular tracking to change as a function of speech intelligibility. In contrast to the modulation observed in neural speech tracking, ocular tracking did not differ between speech intelligibility levels. Our results show the expected differences of higher neural tracking with higher intelligibility, confirming previous findings and speaking in favor of our analysis approach [15, 50]. In general, acoustically intact speech elicits stronger neural tracking and leads to shifts in temporal and spectral signal characteristics [15, 50]. In combination with ocular tracking, a negative relationship between semantic comprehension and ocular speech tracking has been demonstrated [51]. Additionally, the modulation of ocular tracking in response to speech intelligibility has so far only been investigated using a multispeaker experimental design [20], which elicited higher ocular speech tracking of the attended speech stream compared to a single-speaker setting. While the precise relationship between ocular and neural tracking of acoustic signals is not yet fully understood, evidence suggests bidirectional connections between the eyes and the brain [40], which would provide a mechanistic basis for coordinated speech tracking. Applying the logic of these findings to our present results, we propose that neural circuits in relevant processing areas track the higher linguistic structures of speech, while the eyes track attended low-level acoustic features of speech and might track sounds in general, thus responding more to attentional focus rather than intelligibility per se. This enables the integration of the different functional mechanisms of neural and ocular tracking into one coherent framework of speech tracking.

### Increased functional connectivity between eyes and brain sources in late-blind individuals does not lead to enhanced envelope tracking

Sighted participants showed the highest ocular speech tracking values at ∼6 Hz, which intuitively led to the expectation of enhanced connectivity with left-temporal regions at this frequency. Contrary to this assumption, however, late-blind individuals show increased oculo-cerebral coherence compared to both other groups. This increased coupling may reflect enhanced connectivity between auditory and visual systems following late-onset sensory loss. Stronger auditory-to-visual connectivity has previously been reported in late-blind individuals [48], and stronger inter-modal connectivity has been shown in late-deaf individuals with [19] and without [52] cochlear implants. Thus, vision loss at a later stage of development might strengthen oculo-cerebral connectivity. Importantly, this increase does not translate into stronger ocular envelope tracking (see Fig. 1.a.ii) or enhanced neural tracking (see Fig. 1.b.i). While enhanced auditory tracking in occipital regions could, in principle, be driven by eye movements, we did not observe increased occipital speech tracking in the blind groups. Moreover, oculo-cerebral connectivity did not modulate envelope tracking strength. Together, these results suggest that connectivity and tracking reflect independent processes, supporting a distinct interpretation of the connectivity effect. In line with [35], our findings further indicate that oculo-cerebral circuitry is retained in early-blind individuals.

### Limitations

Importantly, current eye-tracking methodologies are optimized for sighted individuals, limiting the precision of gaze-based measures in blind populations. Therefore, we extracted ocular activity using source reconstruction techniques, which is not a well-established eye-tracking method. Nevertheless, we assume that the distinct spatial characteristics of the eyes allow them to be modeled as independent sources. This is also underlined by the fact that both the group and the condition effect in the neural tracking measure differ from those in the eyes, suggesting different activities. Moreover, we cannot disentangle the direction (horizontal vs. vertical) of the eye movements, nor reliably identify saccades or blinks, as eye-movement patterns in the blind groups showed a high variety. Consequently, we refer to all recorded ocular activity collectively as eye movements. To disentangle this, future research should aim to develop improved tracking techniques tailored to blind individuals, extracting saccades and blinks [25] or infrared eye-tracking methods also working with closed eyes [4].

## Conclusion

Our findings provide the first evidence that the oculomotor system tracks auditory information without visual input. Low-frequency (∼1 Hz) ocular activity aligned with the speech envelope across all groups, indicates a hard-wired mechanism supporting auditory processing. In contrast, high-frequency (∼6 Hz) tracking emerged only in sighted individuals, suggesting an experience-dependent process shaped by audiovisual integration. Unlike neural speech tracking, ocular tracking did not vary with intelligibility, implying distinct yet coordinated mechanisms between ocular and cortical systems. Enhanced oculo-cerebral coupling in late-blind individuals further supports adaptive reorganization following sensory loss. Together, these results expand current models of active sensing by demonstrating that eye movements - independent of vision - contribute to the temporal organization of auditory processing, positioning the eyes as active agents in the perception of speech.

## Code Availability

https://gitlab.com/KajaRosa/kbblindaudio

## Data Availability

The authors acknowledge the Austrian NeuroCloud (https://anc.plus.ac.at/), hosted by the University of Salzburg and funded by the Federal Ministry of Education, Science and Research (BMBWF), for providing a FAIR-compliant research data repository. Find the data on request on the https://bids-datasets.data-pages.anc.plus.ac.at/auditory/mablindaudio/

## Author Contributions

Larissa Reitinger wrote the manuscript, gave methodological advice, reviewed the figures and the data analysis, Kaja Benz analyzed the data, Anne Hauswald gave conceptual and methodological supervision and edited the manuscript. Fabian Schmidt gave methodological advice and shared code, Davide Bottari edited the manuscript, Olivier Collignon designed the experiment and edited the manuscript, Nathan Weisz led the data analysis, acquired funding, and edited the manuscript.

## Acknowledgements

We acknowledge our whole research team that supported this work. Markus van Ackeren designed the experiment and recruited the data. The authors acknowledge the computational resources and services provided by Salzburg Collaborative Computing (SCC), funded by the Federal Ministry of Education, Science and Research (BMBWF) and the State of Salzburg. This research was funded in whole or in part by the Austrian Science Fund (FWF) 10.55776/PAT4466224. For open-access purposes, the author has applied a CC BY public copyright license to any author-accepted manuscript version arising from this submission. Markus van Ackeren and Olivier Collignon were funded by the H2020 European Research Council (337573), Kaja Benz was supported by the ÖAW P26-AW2605-P (Austrian Academy of Sciences), Larissa Reitinger was supported by the Austrian Science Fund (FWF; 10.55776/PAT4466224; ”Ocular Speech Tracking”).

## 1 Supplementals

### 1.1 Sanity Checks and mTRF results

**Figure 3:**
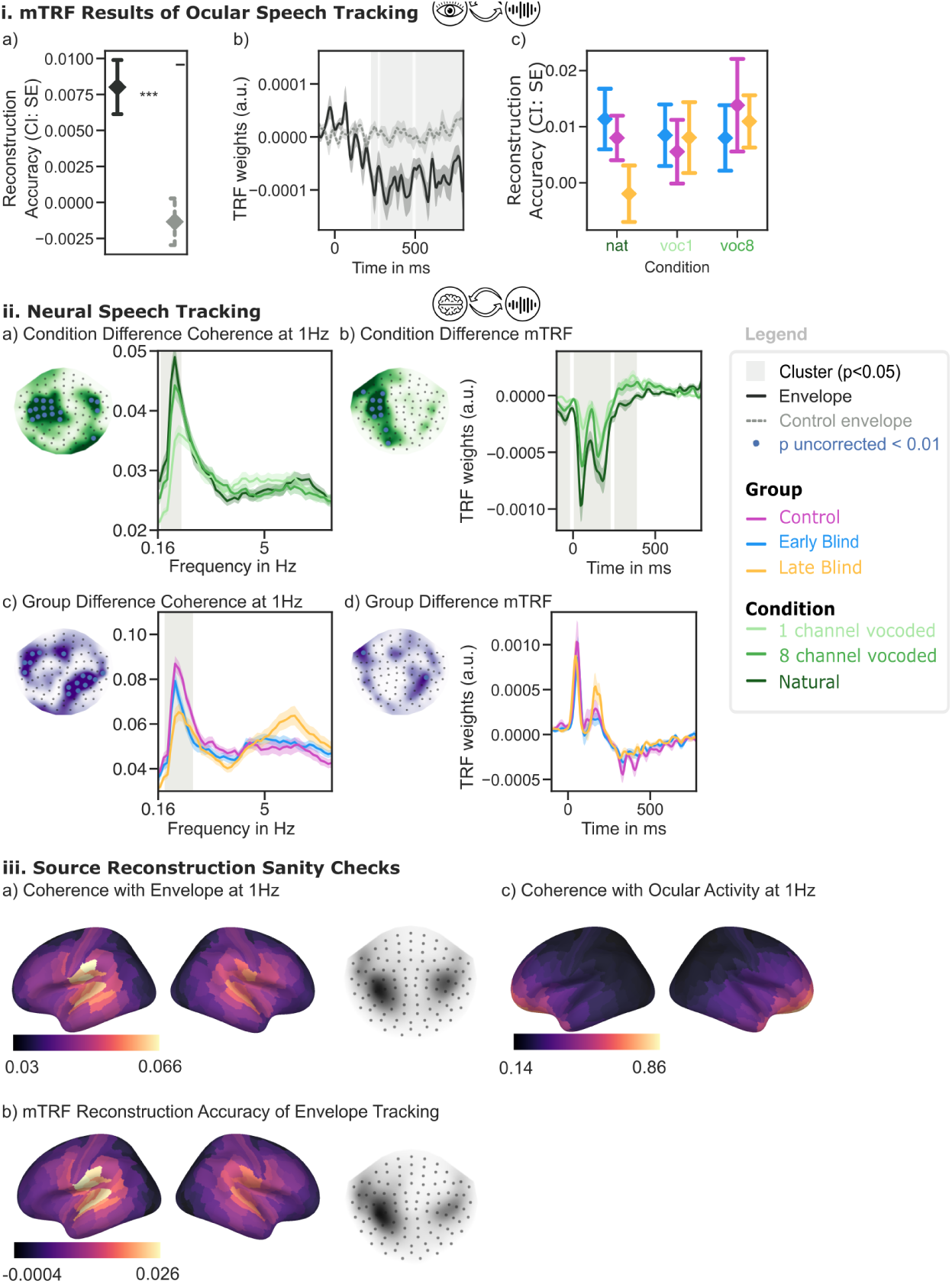
i. Ocular tracking mTRF results. Note that the direction of the mTRFs is not interpretable, direction of the original signals is not meaningful. c) There was no significant main effect of group (*F* (2, 44) = 0.27, *p* = .765, *η*_*g*_^2^ = 0.006), no significant main effect of condition (*F* (44) = 0.73, *p* = .485, *η*_*g*_^2^ = 0.009), and no significant interaction between group and condition (*F* (4, 88) = 0.93, *p* = .448, *η*_*g*_^2^ = 0.022). ii. Condition and group differences of the sensor data for coherence and mTRF. To show maximal differences between groups and conditions, and to make sure that the ocular effects are not based on neural effects we plot the sensors with uncorrected *p <* 0.01 in a time or frequency resolution. iii. Sanity checks of source reconstruction, speech tracking with a) coherence and b) mTRF calculation. c) Coherence of eye movements to neural data at ∼1 Hz.

### Group difference oculo-cerebro connectivity

Cluster from Fig.2.c.ii. Left hemisphere, it starts at 0.167 Hz and ends at 3.33 Hz (*F* (44, 2) = 3187.8, *p <* 0.001) including many regions peaking at the left secondary auditory cortex (L-PBelt-ROI-lh).

On the right hemisphere the cluster peaks occipitaly at the V7 (R-V7-ROI-rh) the significant difference is a bit slower: from 2 until 3.6 Hz (*F* (44, 2) = 608.07, *p <* 0.01).

**Table 1:** Region Names Grouped by F-Value

| F-Sum | Region Names (F-Value Range) |
| --- | --- |
| 30-35 | R-V1-ROI-rh, R-V8-ROI-rh, R-V2-ROI-rh |
| 25-30 | R-V6A-ROI-rh, R-V6-ROI-rh, R-STV-ROI-rh |
| 20-25 | R-V4-ROI-rh, R-V3-ROI-rh, R-PHA1-ROI-rh, R-PreS-ROI-rh |
| 15-20 | R-PGi-ROI-rh, R-ProS-ROI-rh, R-H-ROI-rh, R-VMV2-ROI-rh, R-PFop-ROI-rh |
| 10-15 | R-VMV3-ROI-rh, R-PHA2-ROI-rh, R-PSL-ROI-rh, R-VMV1-ROI-rh, R-DVT-ROI-rh, R-PHA3-ROI-rh, R-PFm-ROI-rh, R-PIT-ROI-rh, R-VVC-ROI-rh, R-IPS1-ROI-rh, R-TPOJ2-ROI-rh |
| 5-10 | R-FFC-ROI-rh, R-MIP-ROI-rh, R-PFcm-ROI-rh, R-TPOJ1-ROI-rh, R-PH-ROI-rh, R-IP2-ROI-rh, R-LO1-ROI-rh, R-PHT-ROI-rh, R-TE2p-ROI-rh, R-LO2-ROI-rh |
| 0-5 | R-V3CD-ROI-rh, R-A5-ROI-rh |

**Table 2:**
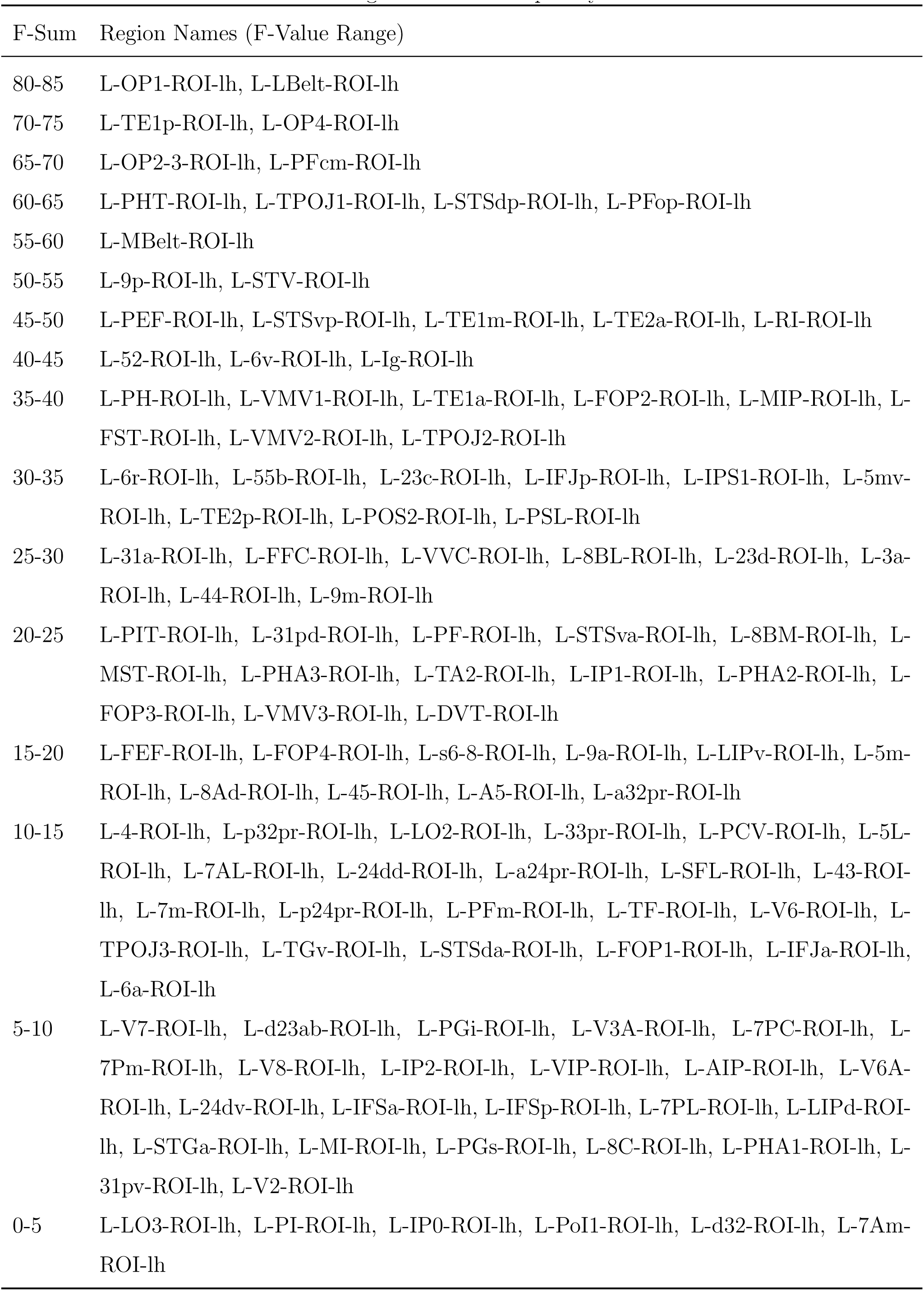
Region Names Grouped by F-Value

